# Microtubules Gate Tau Condensation to Spatially Regulate Microtubule Functions

**DOI:** 10.1101/423376

**Authors:** Ruensern Tan, Aileen J. Lam, Tracy Tan, Jisoo Han, Dan W. Nowakowski, Michael Vershinin, Sergi Simo, Kassandra M. Ori-McKenney, Richard J. McKenney

## Abstract

Tau is an abundant microtubule-associated protein in neurons. Tau aggregation into insoluble fibrils is a hallmark of Alzheimer’s disease and other dementias, yet the physiological state of tau molecules within cells remains unclear. Using single molecule imaging, we directly observe that the microtubule lattice regulates reversible tau self-association, leading to dynamic condensation of tau molecules on the microtubule surface. Tau condensates form selectively permissible barriers, spatially regulating the activity of MT severing enzymes and the movement of molecular motors through their boundaries. We propose that reversible self-association of tau molecules, controlled by the microtubule, is an important mechanism of tau’s biological functions, and that oligomerization of tau is a common property shared between the physiological and disease forms of the molecule.

**One Sentence Summary:** **Reversible tau oligomerization regulates microtubule functions**.

---

In cells, microtubules (MTs) serve as polarized platforms for motor and non-motor proteins. In Alzheimer’s disease, the abundant neuronal MT-associated protein (MAP), tau (*MAPT*), forms insoluble neurofibrillary tangles (NFTs) through aberrant self-association, a process associated with neuronal cell death (*1, 2*). While tau self-association drives NFT formation in disease, a physiological role for homotypic interactions between tau molecules is less clear. Biochemical studies have suggested that tubulin, or MTs, may mediate tau oligomerization (*3*). Additionally, multivalent interactions between unstructured domains can drive liquid-liquid phase separation of tau, promoting tau aggregation (*4*-*7*). When bound to MTs, single molecule studies revealed that tau molecules can exist in either static or diffusive populations, and that tau can bind heterogeneously to MTs (*8*-*10*). While these studies highlight tau’s diverse molecular behaviors, how such behaviors relate to tau’s normal physiological roles in the cell is unknown. Non-aggregated, MT-bound tau has been proposed to regulate the movement of motor proteins and the activity of MT severing enzymes (*9*, *11*-*15*), but a molecular mechanism for how such diverse regulation is achieved is lacking. Using single molecule imaging, here we report that MTs directly regulate the formation of spatially localized tau oligomerization that partitions the MT into distinct domains to regulate diverse biological functions.

We directly observed the binding of recombinantly expressed, GFP-tagged full-length (2N4R) human tau (fig. S1A) to taxol-stabilized MTs *in vitro*. Tau molecules initially bound diffusely along the entire MT lattice, followed by the nucleation and expansion of denser regions of molecules that we term “condensates” due to the localized increase in protein concentration (Fig. 1A, B, and movie S1). Tau condensates expanded along the MT and merged with nearby condensates before reaching a concentration-dependent steady state frequency along the MT lattice, coverage of the MT lattice, and fluorescence intensity (Fig. 1A-D and fig. S1B, C). Condensation was reversed upon removal of soluble tau from solution (Fig. 1B and fig. S1B), suggesting that condensation is distinct from irreversible tau aggregation observed in human tauopathies. We also observed that tau condensation invariably occurred at regions of high MT curvature (Fig. 1E). While tau bound diffusely to MTs assembled with either taxol or the non-hydrolyzable GTP analogue guanosine-5’-[(α,β)-methyleno]triphosphate (GMP-CPP), tau condensates only formed on taxol-, or native GDP MT lattices (Fig 1F-H), revealing that tau condensation is gated by the nucleotide state of the MT lattice. Tubulin’s unstructured C-terminal tails have been reported to affect tau binding and diffusion along MTs (*10*). In our assays, complete removal of tubulin’s tails by subtilisin digestion revealed that tau bound more tightly to digested MTs, but we did not observe tau condensation (Fig. 1I). Conversely, at higher tau concentrations, total tau intensity was greater on undigested MTs suggesting that tau condensation on undigested MTs drives the higher overall affinity of tau for these MTs, as previously observed in bulk biochemical assays (*16*). These results highlight diverse modes of tau’s interaction with MTs, and reveal that tau condensation is dictated by the local curvature, nucleotide state, and presence of the C-terminal tails of tubulin.

**Figure 1:**
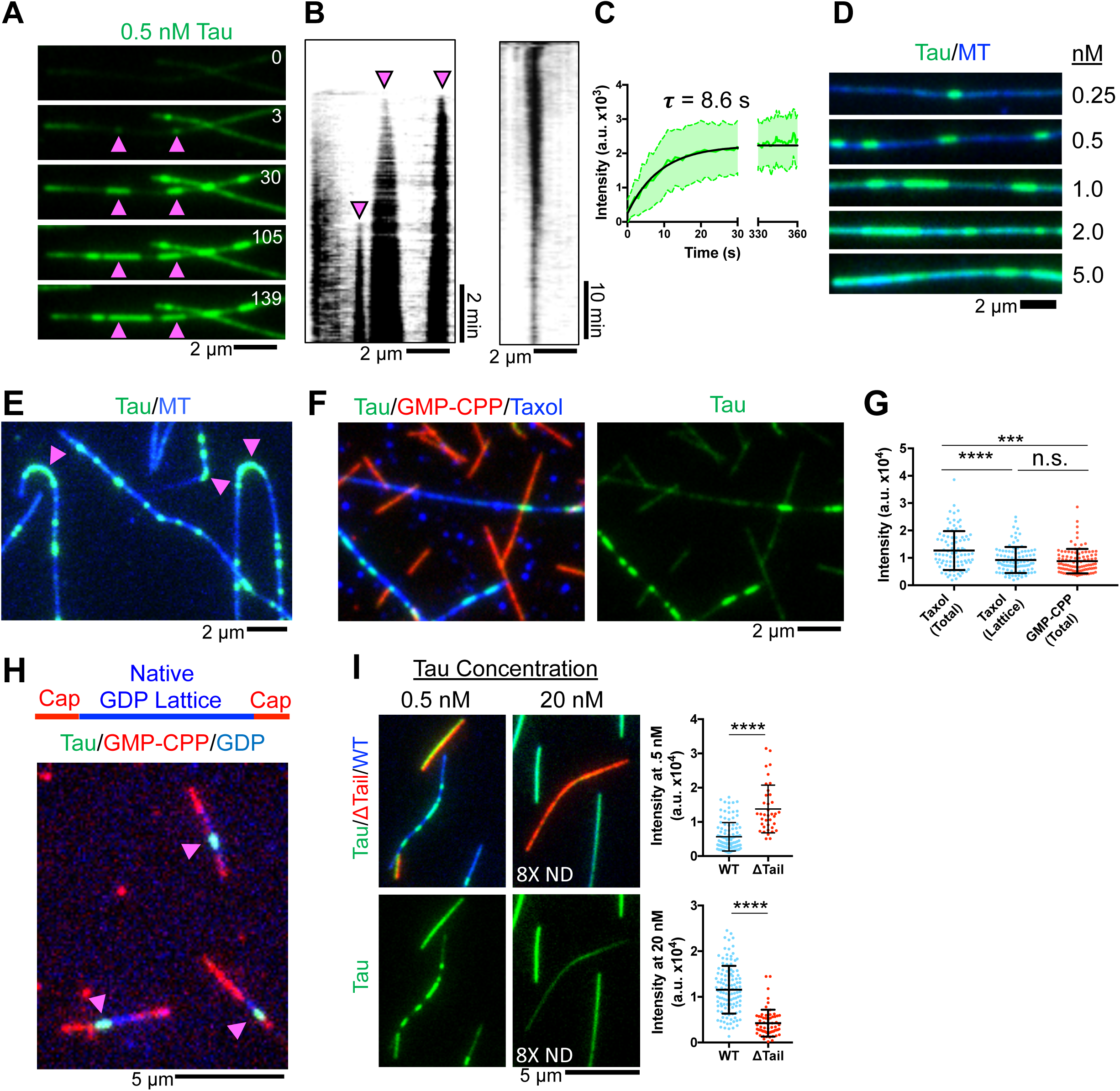
Microtubules Gate the Spatial Condensation of Tau on the Lattice. (A) Time-lapse frames of GFP-tau condensates nucleating and growing on a MT. Pink arrows indicate sites of nucleation. Time in sec. (B) Left: Kymograph of GFP-tau condensates nucleating (magenta arrows) and expanding. Right: Kymograph of a GFP-tau condensate dissolving after tau washout. (C) Quantification of GFP intensity overtime at a condensate nucleation site. (D) Images of MTs at with increasing GFP-tau concentrations, set to equal brightness and contrast. (E) Image of GFP-tau condensates on curved portions microtubules (magenta arrows). (F) Left: Images of 0.5 nM GFP-tau on taxol and GMP-CPP-stabilized MTs. Right: Isolated GFP-tau channel. (G) Quantification of GFP-tau intensity on taxol-stabilized versus GMP-CPP-stabilized MTs. Note the comparison between total intensity including tau condensates (total), and intensity outside of condensates (lattice). (H) Image of 0.5 nM GFP-tau condensates (green) on native GDP MT lattice (blue), stabilized at both ends with GMP-CPP caps (red). Magenta arrows indicate tau condensates. (I) Left: Images of GFP-tau on subtilisin treated (red) and untreated MTs (blue). Below: Isolated GFP-Tau channel. Right: Quantification of GFP-tau intensity. Note the use of an 8X neutral-density (ND) filter in 20 nM tau condition reduces total brightness. ^∗∗∗^P < 0.001, ^^∗∗∗^∗^P < 0.0001, Student’s T-test, one-way ANOVA for multiple comparisons.

Prior experiments have suggested that MTs may mediate the oligomerization of tau (*3*), but direct observation of this phenomenon is lacking. We hypothesized that tau condensates may be a novel form of homotypic tau interactions, and set out to determine the biophysical nature of tau condensation. Fluorescence recovery after photobleaching (FRAP) experiments revealed that tau condensates recovered ~ three-fold slower than diffusely bound tau outside of condensate boundaries (Fig. 2A). To directly observe tau dynamics within condensates, we utilized a single-molecule spiking assay in which SNAP-TMR-labeled tau was added at a 1:40 ratio of GFP-labeled tau (Fig. 2B, C, and movie S2). Outside of condensates, single SNAP-TMR tau molecules rapidly diffused along the MT lattice with an average dwell time of 6.2 s (Fig. 2B, C). Single tau molecules diffused to condensate boundaries upon which their behavior altered dramatically, transitioning from rapidly diffusing to statically bound (Fig. 2B), suggesting that interactions with GFP-tau molecules that compose the condensate dramatically reduced the molecular dynamics of SNAP-TMR tau on the MT lattice. Within the condensates, dwell times for single tau molecules increased six-fold, indicating cooperative interactions between tau molecules drive condensate formation and maintenance (Fig. 2B, C). In further support of the dynamic nature of tau condensates (Fig. 1B-C), we observed single tau molecules could transition into and out of the condensate boundary, switching behavior between immobility and rapid diffusion repeatedly (Fig. 2B). These results reveal that cooperative interactions between tau molecules, gated by the MT lattice (Fig. 1), strongly reduce tau’s molecular dynamics, providing a possible molecular mechanism for localized tau condensation.

**Figure 2:**
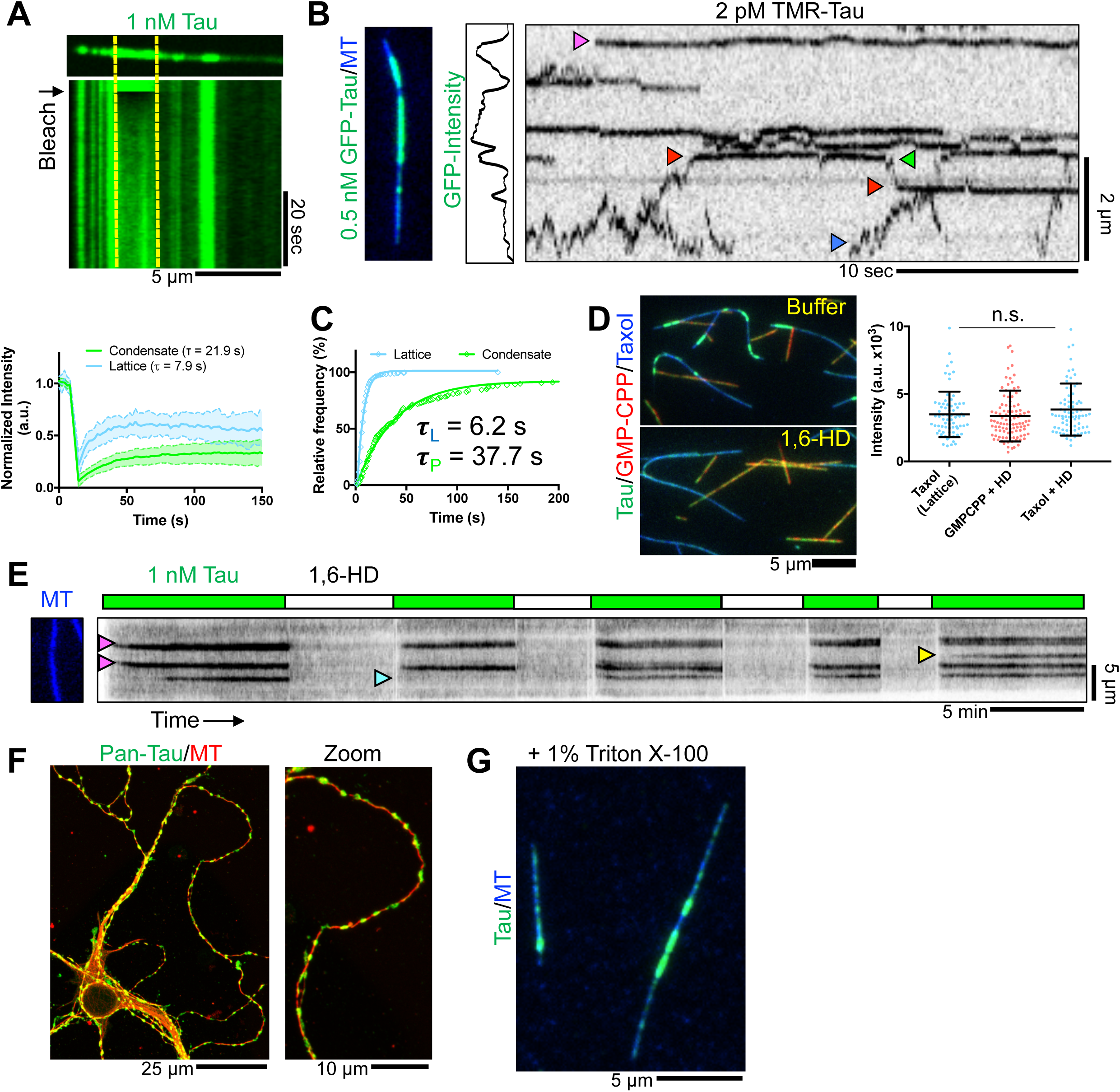
Tau Condensation is Driven Hydrophobic Interactions at Specific Nucleation Sites on the MT Lattice. (A) Above: Image and kymograph of a MT with GFP-tau condensate undergoing FRAP. Yellow lines represent photobleached region. Below: Quantification of FRAP recover in both condensate and diffusely decorated MT lattice regions, with calculated recovery constants. (B). Image of a MT with a GFP-tau condensate, GFP intensity plot along this MT, and kymograph of SNAP-TMR-labeled tau molecules. Magenta arrow indicates a static tau molecule within a condensate. Blue arrow indicates a diffusive tau outside of the condensate. Red arrows denote events where a molecule enters, and green arrows denote exits events from a condensate. (C) Cumulative frequency plot of SNAP-TMR-tau dwell times either within GFP-tau condensates or on the MT lattice outside of condensates. (D) Left: Images of 0.5 nM GFP-tau bound to taxol or GMP-CPP stabilized MTs in the presence of buffer or buffer with 10% 1,6-HD. Note the disappearance of tau condensates in the presence of 1,6-HD. Tau concentration was kept constant during buffer exchange. Right: Plot of GFP intensity outside of condensates (lattice) or total GFP intensity in the presence of 1,6-HD. (E) Kymograph of alternating washes of 1 nM GFP-tau with or without 8% 1,6-HD. Washing scheme diagrammed above. Magenta arrows denote condensate nucleation events, blue denotes a condensate that failed to reform once after 1,6-HD washout, yellow denotes a condensate that formed subsequent to initial tau introduction. (F) Images of immunostained DIV7 mouse hippocampal neurons. (G) Image of *in vitro* GFP-tau condensates on MTs in the presence of 1% Triton X-100. Student’s T-test, one-way ANOVA for multiple comparisons.

Tau condensates are reminiscent of reversible membraneless liquid-liquid phase separations (LLPS) previously observed for tau in solution (*5*, *17*). Therefore, we exposed tau condensates to 1,6-hexanediol (1,6-HD), an aliphatic alcohol shown previously to dissolve tau LLPS droplets (*17*). Surprisingly, tau condensates were completely dissolved by 1,6-HD, without affecting diffuse binding to either taxol or GMP-CPP MTs (Fig. 2D). Thus, hydrophobic interactions between tau molecules are necessary for condensation, but not MT binding. We used 1,6-HD to further probe the role of the MT lattice in condensation. Tau condensates formed, dissolved, and re-formed largely at the same locations along the MT lattice, and we infrequently observed the formation of a new condensate, or the lack of condensate reformation after 1,6-HD removal (Fig. 2E and movie S3). This observation strongly indicates that local regions of the MT lattice act as nucleation ‘hot-spots’ for tau condensation. We hypothesize that these hot-spots represent local areas of lattice distortion, similar to the highly curved regions that invariably nucleate tau condensates (Fig. 1E). Our results thus far show that tau condensation is reversible, and driven by the same types of interactions between tau molecules that lead to LLPS of tau in solution. The slow recovery of condensates after FRAP differs from the typical properties of a solution phase LLPS system (*17*). Tau condensates are kinetically more stable, which we hypothesize may be due to scaffolded interactions with the MT. We note that tau condensation occurs in physiological buffer, at concentrations an order of magnitude below those shown previously for tau in non-physiological solutions (*5*, *7*, *17*), which we hypothesize may be due to locally high concentrations of tau molecules bound to the MT lattice.

We next sought evidence that tau condensation can occur *in vivo*. We stained mouse hippocampal neurons with two different pan-tau antibodies and observed developmentally dependent tau localization to focal puncta along MTs, resembling *in vitro* tau condensates (Fig. 2F, and fig. S2). Similar staining has been reported in cultured neurons (*18*-*20*), with some suggesting that because such puncta are resistant to Triton X-100 extraction, they represent tau irreversible aggregates (*21*). However, *in vitro* assembled tau condensates were similarly resistant to Triton X-100 (Fig. 2G), calling into question this interpretation. Thus, tau localization in neurons is diverse, and focal puncta of tau resembling tau condensates form inside of neurons when neurite maturation reaches a certain threshold.

Next, we set out to determine what domains mediate tau-tau interactions within condensates. In neurons, six tau isoforms exist with differences in the number of projection domain inserts (N) and number of MT binding repeats (R). All isoforms contain a microtubule binding domain (MTBD) flanked by a proline-rich region and pseudo-repeat region (Fig. 3A). Alternative splicing in the MTBD or projection domain insert region did not grossly perturb tau condensation (Fig. 3A, B). We assayed for tau oligomerization by first forming condensates with full-length mScarlet-tagged tau (2N4R), followed by introduction of equimolar amounts of GFP-tagged tau isoforms (2N4R/2N3R/0N3R). We measured enrichment of GFP-tau isoform signal within full-length mScarlet-tau condensates versus GFP signal outside of mScarlet-tau condensates. We observed a two- to three-fold enrichment of GFP-2N4R, -2N3R, and -0N3R isoforms within condensates, indicating that alternative splicing does not grossly affect incorporation into 2N4R tau condensates (Fig. 3C, and fig. S3).

**Figure 3:**
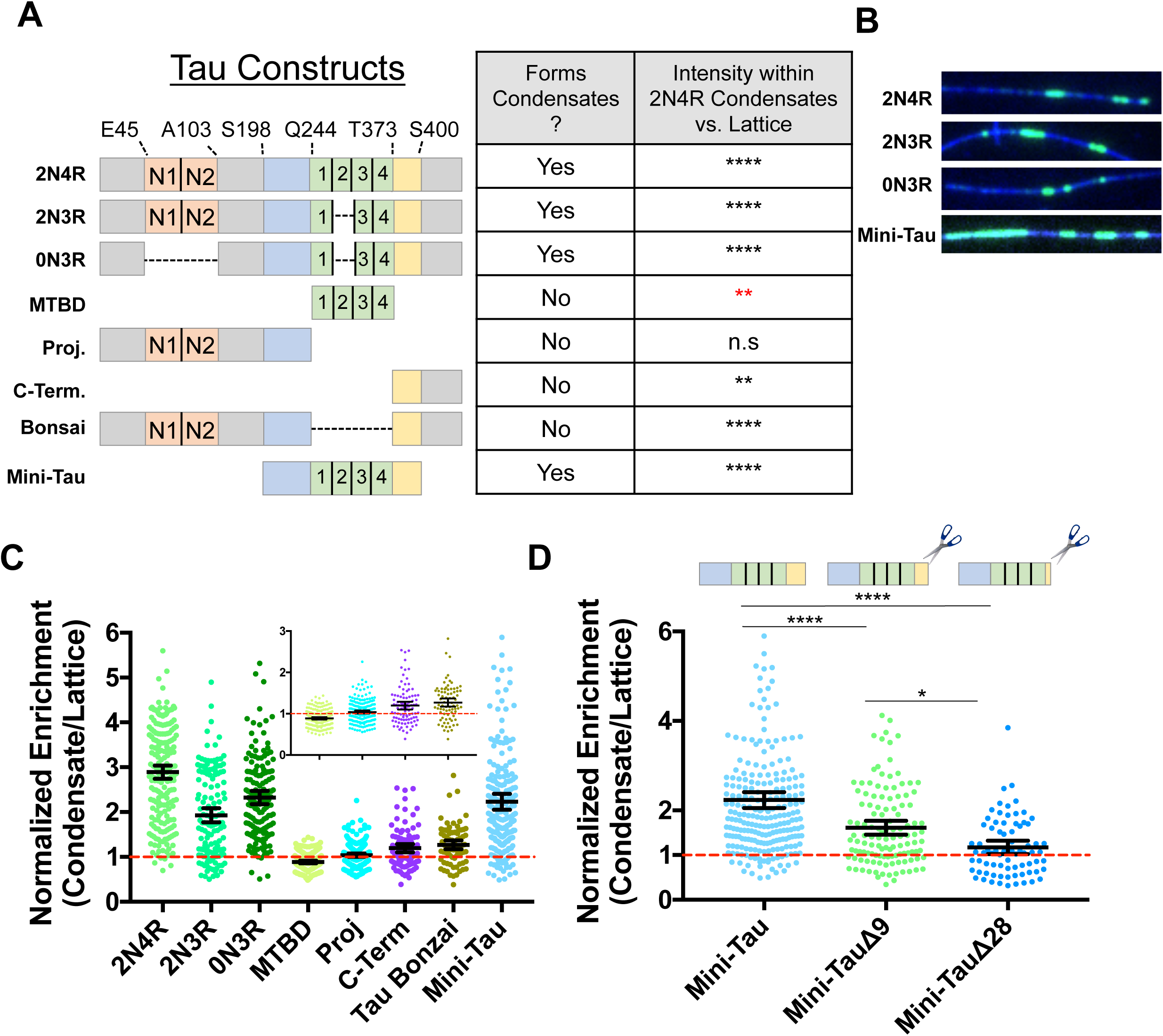
The C-terminal Pseudo-Repeat Region of Tau Licenses the Rest of the Molecule Into Tau Condensates. (A) Schematic of tau isoforms and constructs. Orange boxes: alternatively spliced N-term. inserts. Blue: proline-rich domain. Green: MT binding repeats. Yellow: pseudo-repeat domain. Right, table summarizing ability of different tau constructs to form condensates and statistical significance of quantitation of tau construct intensity within 2N4R condensates versus on MT lattice (see also (fig. S3)). (B) Images of tau condensates formed from different alternatively spliced or artificially truncated tau constructs. (C) Quantification of the fold enrichment of various tau constructs into 2N4R tau condensates versus the MT lattice surrounding the condensate. Inset shows zoom for clarity. Error bars: 95% C.I. (D) Quantification of the fold enrichment of Mini-Tau and C-terminal deletion constructs into 2N4R tau condensates. Data for Mini-Tau reproduced from (C) for comparison. Error bars: 95% C.I. ^∗^P < 0.05, ^∗∗^ P < 0.01, ^∗∗∗^P < 0.001, ^∗∗∗∗^P < 0.0001. Student’s T-test, one-way ANOVA for multiple comparisons.

We truncated domains of tau and found that, in contrast to previously published data on tau LLPS, the MTBD alone was weakly excluded from condensates (Fig. 3A, C, and fig. S3C), while the isolated projection domain exhibited diffuse binding (Fig. 3A, C, and fig. S3C). The isolated C-terminus of tau segregated into condensates, though more weakly than full-length tau (1.2-fold vs. 2.9-fold respectively), as did a “bonsai” construct consisting of a fusion between the N- and C-terminus, but lacking the MTBD (1.3-fold, Fig. 3A, C, and fig. S3C). These data indicate that the C-terminus of tau licenses other portions of the molecule into condensates. Consistently, while the MTBD was de-enriched from condensates, the addition of the flanking N-terminal proline-rich and C-terminal pseudo-repeat domains (mini-tau) restored segregation into condensates to near full-length tau levels (Fig. 3A, C, and fig. S3D). The C-terminal pseudo-repeat domain is evolutionarily conserved and rich with aliphatic hydrophobic residues (fig. S4A). Truncation of residues from the pseudo-repeat region of mini-tau resulted in a progressive decrease in condensate incorporation, further indicating that tau condensation requires interactions located within this region (Fig. 3D, and fig. S3D). However, removal of the entire pseudo-repeat domain (mini-tau Δ28) largely, but not completely, abolished incorporation into 2N4R tau condensates (Fig. 3D, and fig. S3D). Previous studies have implicated the proline-rich region and MTBD in MT binding (*22*), while our mapping data presented here reveals that hydrophobic interactions within the pseudo-repeat region are essential for interactions that drive tau condensation on the MT lattice.

Next, we wondered about the biological effects of tau condensation. Previous studies have suggested that tau can regulate the movement of molecular motors, including the cytoplasmic dynein-dynactin complex (*9*, *23*). However, the reported effects of tau on dynein-dynactin movement are not broadly consistent with the strongly processive movement of activated dynein-dynactin-cargo adapter complexes discovered subsequently (*24*, *25*). We thus focused on the effects of tau condensation on minus-end directed motor transport driven by activated dynein-dynactin motor complexes to reconcile these findings. We found the majority of processive dynein-dynactin-BicD2N (DDB) complexes passed through condensates (83.9%), often displaying dramatic pausing at the condensate border (49.4%) (Fig. 4A, and fig. S5A, B, movie S4). A small population of processive motor complexes (~3%) exhibited unidirectional movement before switching to a bidirectional state at tau condensates (fig. S5B, C). Similarly, a fraction of DDB motors that displayed only diffusive behavior on the MT (*24*) consistently reversed direction at tau condensates, (Fig. 4B, and fig. S5B). This behavior was very similar to a purified p150^*glued*^ construct (Fig. 4B), suggesting that condensates are permissive only to processive dynein movement, but not to dynactin-mediated diffusion. Thus, in contrast to the plus-end directed motor kinesin (*9*, *26*, *27*), processive dynein is physically able to pass through tau condensates. The difference in behaviors between the motors could be explained by recent cryo-EM structures of tau on the MT (*28*). While the kinesin motor domains sterically clashes with MT-bound tau, we found that the much smaller dynein MT binding domain does not (Fig. 4C, and fig. S5D-F).

**Figure 4:**
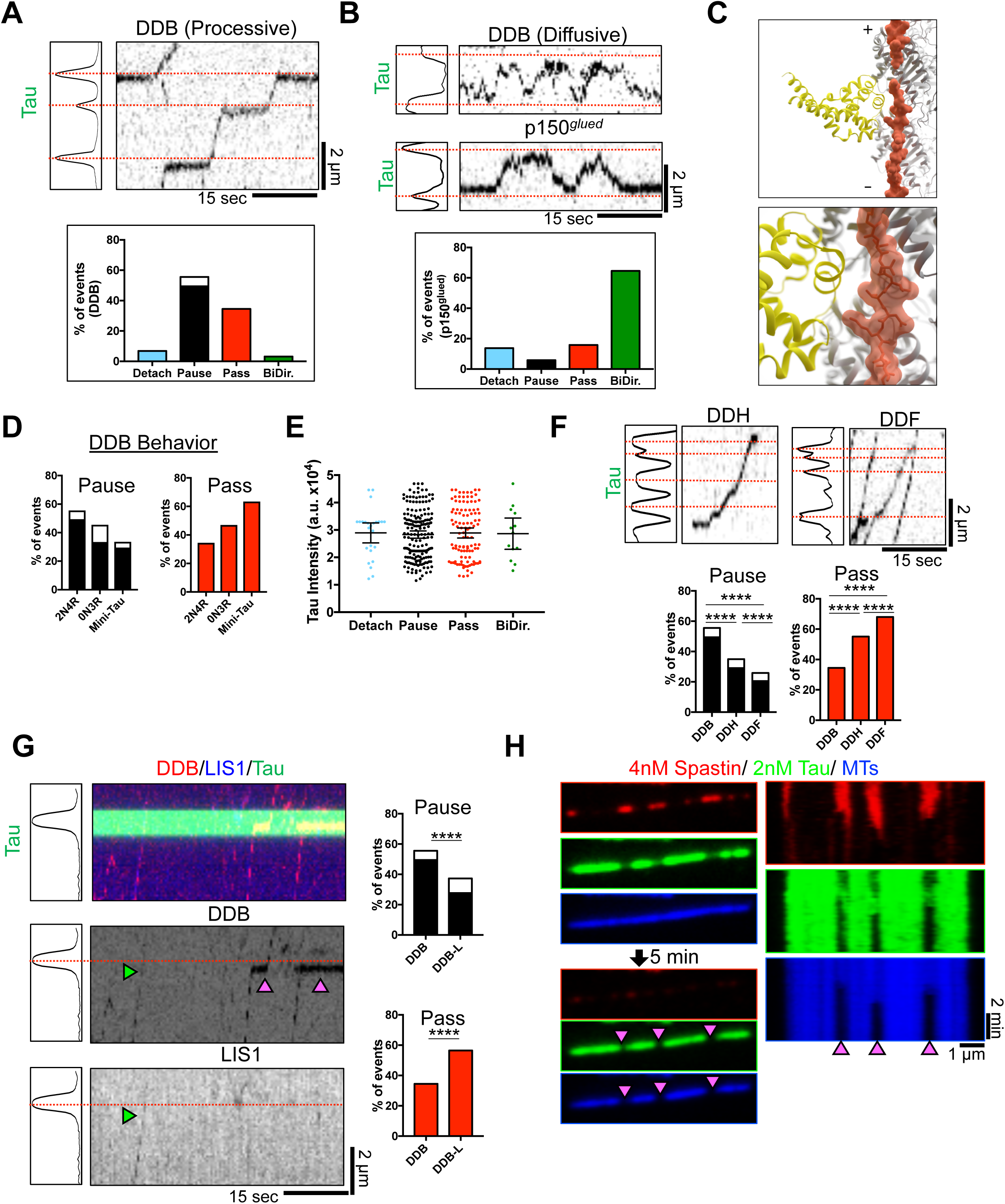
Tau Condensates Form Selectively Permissible Barriers to Regulate Diverse MT Functions. (A) Top: GFP-tau intensity plot of with accompanying kymograph of processive DDB. Bottom: Graph of event distribution for DDB. White box indicates proportion of molecules that detach while paused. (B) Top: Intensity graph of with accompanying kymograph of diffusive DDB and p150^*glued*^. Bottom: Event distribution for p150^*glued*^. (C) Model of tau (R2x4, pdb: 6CVN; orange, (*28*)) and MT-binding footprint the dynein MTBD (*DYNC1H1*, pdb: 3JLT; yellow). (D) Event distribution of pausing and passing DDB behavior at 0N3R and Mini-Tau condensates. Distribution of DDB behavior at 2N4R condensates reproduced from (A) for comparison. (E) Distribution of tau condensate peak intensity for each DDB behavior. (F) Left: Kymograph and accompanying tau intensity of DDH and DDF. Right: Passing and pausing distribution for DDH and DDF. Distribution of DDB behavior reproduced from (A) for comparison. See fig. S5B for full distribution of motor behaviors. (G) Left: Kymograph and accompanying tau intensity of DDB-L. Right: Passing and pausing distribution for DDB-L. Distribution of DDB behavior reproduced from (A) for comparison. See fig. S5B for full distribution of motor behaviors. Green arrow denotes DDB-L complex, magenta arrows DDB complexes. (H) Left: Images of mScarlet-tau condensates in the presence of 4 nM GFP-spastin before and after 5 min. incubation. Right: Kymographs from spastin, tau, and MT channels. magenta arrows denote regions of spastin-mediated MT destruction. ^∗^P < 0.05, ^∗∗^P < 0.01, ^∗∗∗^P < 0.001, ^∗∗∗∗^P < 0.0001. Student’s T-test, one-way ANOVA for multiple comparisons.

We sought to determine the domains of tau that contribute to the dramatic pausing of processive dynein complexes (Fig. 4A). Condensates formed from the shortest natural tau isoform (0N3R) or the mini-tau construct, allowed progressively greater numbers of processive motor complexes to pass unimpeded (47% and 63.4%, respectively) (Fig. 4D, and fig. S5B). Further, we found that tau density within the condensate, was not a predictive determinant of motor behavior upon encountering condensates (Fig. 4E), indicating that the distribution of motor behaviors was stochastic, rather than dependent upon tau density

Recent biophysical and structural data have defined adaptor-dependent variations in the number of dynein dimers linked to the dynactin scaffold (*29*, *30*). BicD2N has been shown to favor only one dynein dimer, while the adapter Hook3 largely recruits two dynein dimers per dynactin. Dynein-dynactin-Hook3 (DDH) complexes passed through tau condensates without pausing at significantly higher rates (55%) compared to DDB (34.5%), indicating that two scaffolded dynein dimers are better able to navigate tau condensates unimpeded (Fig. 4F, and fig. S5B). Another complex, dynein-dynactin-FIP3 (DDF), exhibited significantly increased rates of passing through tau condensates without pausing (68%), indicating that FIP3, likely recruits a pair of dynein dimers similar to Hook3 (Fig. 4F, and fig. S5B). Furthermore, the dynein regulatory protein LIS1 directly impinges on dynein’s mechanochemistry, and allosterically controls DDB velocity (*31*-*34*). Analysis of DDB complexes bound to Lis1 (DDB-L) revealed that these complexes were also better able to navigate tau condensates unimpeded (58.6% vs. 34.5%) (Fig. 4G, and fig. S5B). These results reveal that allosteric control of dynein motor activity through multiple mechanisms regulates the motor’s ability to pass through tau condensates unimpeded, and suggest that tau condensates may regulate the movement of retrograde traffic in a cargo-dependent manner.

Finally, we explored how non-motor MAPs were affected by tau condensates. The molecular mechanism for tau-mediated inhibition of MT severing is unknown. We found that a truncated, active form of the MT severing enzyme spastin (*35*) was largely excluded from tau condensates (Fig. 4H). As a result, tau condensates protected the underlying MT lattice from spastin-mediated severing, while the lattice surrounding the condensate was destroyed (Fig. 4H and movie S5). These results demonstrate that tau condensates regulate diverse MT-based functions by acting as selectively permissible barriers for MT effector proteins.

In summary, our work has uncovered a novel, regulated form of reversible tau oligomerization that partitions the MT lattice into functional subdomains. Complementary work presented in Siahaan et al. confirms this behavior of tau and extends our observations of how reversible tau oligomerization regulates molecular motor transport and microtubule turnover. We propose that tau condensation represents a physiological form of tau self-association, regulated and scaffolded by the MT lattice, which can be harnessed by cells to spatially direct diverse MT-based molecular pathways. Further, our results reveal that reversible oligomerization allows tau to perform biologically meaningful functions, demonstrating that oligomerization is important in both physiological and pathological roles of tau. Because tau condensation is sensitive to overall tau concentration, we speculate that loss of tau monomer to alternative self-association pathways, such as fibrillization, will negatively impact tau condensate formation, maintenance, and function in cells.

## Acknowledgments

We would like to thank all the members of the McKenney and Ori-McKenney labs for critical feedback, as well as Jawdat Al-Bassam, Marcus Braun, and Stefan Diez for feedback and the sharing of data:

## Author contributions

R.J.M. and R.T., and K.M.O.M. conceived of the project. R.T., A.L. and T.T. produced reagents. R.T. performed all in vitro experiments. J.H. and S.S. provided hippocampal neuron cultures, D.W.N. created molecular models, M.V. assisted with data analysis;

## Data and materials availability

All data is available in the main text or the supplementary materials.

## Supplementary Materials

### Materials and Methods

#### Microtubule Assembly

Porcine brain tubulin was isolated using the high-molarity PIPES procedure as described and then labeled with biotin-, Dylight-405 NHS-ester, or Alexa647 NHS-ester as described (http://mitchison.hms.harvard.edu/files/mitchisonlab/files/labeling_tubulin_and_quantifying_labeling_stoichiometry.pdf). Microtubules were prepared by incubation of 100 uM tubulin with 1mM GTP for 10 min. at 37°C, followed by dilution into 20 μM final taxol for an additional 20 min. GMP-CPP MTs were prepared similarly but with 1mM GMP-CPP instead of GTP without taxol. Microtubules were pelleted at 80K rpm over a 25% sucrose cushion in a TLA-100 rotor and the pellet was resuspended in 50 μL BRB80 containing 10 μM taxol. For removal of tubulin C-terminal tails, microtubules were further treated with subtilisin for 1 hour at 37 degrees as described *(23)*. The reaction was terminated by 1 mM PMSF and pelleted at 80K rpm as before. Concentration of subtilisin used and digestion were assayed by Coomassie staining and recombinant p150^glued^ binding (*36*). GMP-CPP capped microtubules were prepared as previously described by (*37*).

#### Protein Constructs

All human tau and spastin constructs were cloned into pET28A vector using Gibson assembly. Constructs contain a N-terminal cassette consisting of: a 6x His-tag and tandem Strep-tags connected by a GS-linker. The sequence is as follows: MGSSHHHHHHSSGLVPRGSHMWSHPQFEKGGGSGGGSGGSAWSHPQFEKGS. This cassette is then followed by the fluorophore (sfGFP/mScarlet/SNAPf) followed by a precision protease cleavage site. Human spastin cDNA was purchased from Transomics (BC150260). A fully active, truncated spastin (Δ227) *(34)* was cloned into pET28-strepII-sfGFP. Full-Length human tau was purchased from Addgene (#16316). Amino acid boundaries for tau constructs are as described in Figure 3A. In short, the projection domain inserts were from E45-T102. The Proline-rich domain encompassed S198-L243, and the MTBD was defined as Q244-E372. The second repeat (exon 10) removed in 3R tau constructs spans K274-G304. The pseudo repeat region consists of T373-V399.

#### Protein Purification

Tau and Spastin were expressed in BL21(DE3) cells (Agilent). The cells were grown at 36° C until OD_600_ of 0.6, then induced with 0.4 mM IPTG overnight at 18 ° C. Cells were resuspended in buffer X and lysed using an Emulsiflex C-3 (Avestin). Proteins were affinity-purified on Strep XT beads (IBA). Tau constructs were further purified by anion exchange on a HiTrap Q HP column in Protein Buffer pH 7.5 (50 mM Tris-HCl pH 7.5, 2 mM MgCl_2_, 1mM EGTA, and 10% glycerol) with a salt gradient from 100 mM to 400 mM. Full-length tau constructs were further purified by size exclusion chromatography on a Superose 6 column in Protein Buffer pH 8. All Mini-Tau based constructs were induced for only 4 hours and were purified similarly to all other tau construct. For Mini-Tau constructs and Spastin, we performed cation exchange on a HiTrap SP HP column with the same conditions as other tau constructs. Dynein-Dynactin-Cargo Adaptor complexes were purified from rat brain lysate as described *(23)*. Briefly, all SNAPf-tagged adapter protein constructs were purified by Strep-tag affinity as above and further purified by size exclusion chromatography on a Superose 6 column in 60 mM Hepes pH 7.4, 50 mM K-acetate, 2 mM MgCl_2_, 1 mM EGTA, 10 % glycerol. Dynein-Dynactin-adapter complexes were labeled with in a ~4:1 ratio of dye:SNAPf-tagged protein at 2 μM SNAP-TMR, SNAP-Alexa647, or SNAP-Alexa488 dye (NEB) during the isolation procedure and were frozen in small aliquots and stored at -80°C. The protein concentration was assessed using a Nanodrop One (ThermoFisher). Protein concentrations given are for the total amount of fluorophore (monomer) in the assay chamber. All buffers and chemicals were from Sigma Aldrich.

#### TIRF Microscopy

All TIRF microscopy was performed on a custom built through the objective TIRF microscope (Technical Instruments, Burlingame CA) based on a Nikon Ti-E stand, motorized ASI stage, quad-band filter cube (Chroma), Andor laser launch (100 mW 405 nm, 150 mW 488 nm, 100 mW 560 nm, 100 mW 642 nm), EMCCD camera (iXon Ultra 897), and high-speed filter wheel (Finger Lakes Instruments). All imaging was performed using a 100X 1.45NA objective (Nikon) and the 1.5X tube lens setting on the Ti-E. Experiments were conducted at room temperature. The microscope was controlled with Micro-manager software (*38*). For imaging Tau binding at 20 nM (Fig. 1I), an 8X neutral density filter was used to reduce total signal intensity.

TIRF chambers were assembled from acid washed coverslips (http://labs.bio.unc.edu/Salmon/protocolscoverslippreps.html) and double-sided sticky tape. Taxol-stabilized MTs were assembled with incorporation of ~ 10% Dylight-405- or Alexa647-, and biotin-labeled tubulin. Chambers were first incubated with 0.5 mg/mL PLL-PEG-Biotin (Surface Solutions Inc.) for 10 min., followed by 0.5 mg/mL streptavidin for 5 min. Microtubules were diluted into BC Buffer (80mM Pipes pH 6.8, 1mM MgCl_2_, 1mM EGTA, 1 mg/mL BSA, 1mg/mL casein, 10μM taxol) then incubated in the chamber and allowed to adhere to the streptavidin-coated surface for 10 minutes. Unbound MTs were washed away with TIRF buffer (60 mM Hepes pH 7.4, 50 mM K-acetate, 2 mM MgCl_2_, 1 mM EGTA, 10 % glycerol, 0.5 % Pluronic F-127, 0.1 mg/mL Biotin-BSA, 0.2 mg/mL κ-casein, 10μM taxol). Unless otherwise stated, experiments were conducted in imaging buffer (60 mM Hepes pH 7.4, 50 mM K-acetate, 2 mM MgCl_2_, 1 mM EGTA, 10 % glycerol, 0.5 % Pluronic F-127, 0.1 mg/mL Biotin-BSA, 0.2 mg/mL κ-casein, 10μM taxol, 2 mM Trolox, 2 mM protocatechuic acid, ~50 nM protocatechuate-3,4-dioxygenase, and 2 mM ATP) as in (*39*). Unless specifically stated, all tau assays were performed with 0.5 nM tau.

The resulting data was analyzed manually in ImageJ (FIJI). For images displayed in figures, background was subtracted in FIJI using the ‘subtract background’ function with a rolling ball radius of 50 and brightness and contrast settings were modified linearly. In images where there was substantial drift, the “Descriptor-based series registration (2D/3D + T)” plug-in was used in FIJI with interactive brightness and size detections in the MT channel to register the images.

#### Continuous imaging assays

Tau condensation assays (Fig. 1A-C, S1A) were conducted in Cellvis 96-well Glass Bottom Plate (Cellvis, #P96-1.5H-N) as previously described (*39*). For washout experiments (Fig. 1B, S1A, 2DE), double-sided sticky tape chambers were assembled with 22 x 40 mm coverglass perpendicular to the glass slide. The coverslip was then sealed to the coverglass by epoxy. The extra coverglass area allowed for seamless buffer exchange during imaging. For wash-in (Fig. S1A), and 1,6-hexanediole experiments (Fig. 2DE), images were taken one second apart until the end of the assay. For wash-out assays, images were taken every 30 seconds.

Intensity was measured manually using kymographs on ImageJ. Time zero was defined as the point of visible nucleation.

#### Photobleaching Experiments

FRAP experiments: Images were acquired on a Olympus FV1000 confocal microscope with a 63x/1.42 oil immersion lens at 2 seconds per image. All experiments were done at 25°C. For the FRAP experiments a pre-bleach image was acquired by averaging 8-10 consecutive images. Then 10 regions were bleached (2 background, 4 condensate, 4 lattice) at 100% power without scanning. Images were then taken at 2-s intervals. The microscope was controlled by Slidebook6 software.

Plots were generated using Slidebook6 analysis software and exported to excel. The background subtracted average intensity was measured in an region of interest (ROI) overtime and normalized to the initial fluorescence intensity within the first 8 frames. Data from 24 condensate and lattice regions were analyzed and pooled from experiments from 3 days with 2 different protein preparations.

#### Single molecule spiking experiments

2N4R SNAPf-Tau was labeled with TMR dye in a 1:4 molar ratio for 2 hours before size exclusion with Zeba Spin Desalting Column (Thermo Scientific #89882). Then, 0.5 nM GFP-Tau was flowed in to first form condesnates. Next, a mixture of 0.5 nM GFP-Tau and 10-20 pM TMR-Tau was flowed in the chamber. Images in the TMR channel were taken every 0.25 seconds. Images were taken every second to measure single molecule dwell times. We used the GFP channel as a fiducial for dwell times inside and outside of condensates. Only molecules whose entire residency on the MT lattice was captured, inside or outside condensates, were counted.

#### Primary Neuronal Cultures

Cultured neurons were obtained from embryonic day 16.5 (E16.5) mouse embryonic hippocampi. Hippocampi were carefully dissected and meninges removed. Dissociation was achieved by a combination of enzymatic digestion using papain and pipetting homogenization (Worthington Biochemical Corporation). Neurons were resuspended in neuron growth media (Neurobasal media containing 2% B27 supplement, 2% GlutaMAX solution, glucose, and penicillin/streptomycin). A total of ~10^5^ neurons were plated in poly-D-Lysine-coated coverslips. Growth media was changed every two days. The morning a vaginal plug was observed was considered E0.5. All animals were used with approval from the University of California Davis Institutional Animal Care and Use Committees.

At the appropriate day after isolation, the neurons were fixed in 4% PFA for 20 minutes and permeabilized with 0.3% Triton-X for 5 minutes. Neurons were blocked with 5% BSA and incubated with primary antibody (Tau – Genetex #49353, or Thermo Scientific #13-6400, Beta-Tubulin – Abcam #ab6046), followed fluorescently-labelled secondary antibody (Anti-chicken – A11039, Anti-mouse – A28175, A28180, Anti-rabbit - A27039), and mounted with Vectashield.

Coverslips were imaged on a Leica TCS SPE-II RYBV with automated DMi8 with a Leica laser launch (25 mW 405 nm, 10 mW 488 nm, 20 mW 561 nm, 18 mW 635 nm). All imaging was performed using a HC PL APO CS2 63X 1.40NA objective (Leica) Experiments were conducted at room temperature. The microscope was controlled with Leica LAS X software and analyzed with ImageJ.

At the appropriate day after isolation, the neurons were fixed in 4% PFA for 20 minutes and premiabilized with 0.3% Triton-X for 5 minutes. They were then blocked with 5% BSA and incubated with primary antibody. They were then incubated with secondary antibody and mounted with Vectashield. Coverslips were imaged on a Leica TCS SPE-II RYBV with automated DMi8 with a Leica laser launch (25 mW 405 nm, 10 mW 488 nm, 20 mW 561 nm, 18 mW 635 nm). All imaging was performed using a HC PL APO CS2 63X 1.40NA objective (Leica) Experiments were conducted at room temperature. The microscope was controlled with Leica LAS X software and analyzed with ImageJ.

#### Data Analysis for Condesnate Enrichment (Fig. 3)

mScarlet-tagged 2N4R tau condensates were used as fiducials for condensate boundaries. Background subtracted mean intensities were obtained for a linescan along the microtubule. Each straight and uninterrupted (no MT overlaps) stretch of MT was counted as a single data point. Data points from two different protein preparations of mScarlet-labeled 2N4R tau condensates were pooled. Fold enrichment was calculated by dividing each data point for condensate intensity by the average value of associated lattice intensity.

#### Dynein-Dynactin-Adapter Behavior Assays

DDX movement was visualized manually using kymographs as described in (*39*). The behaviors at condensates were characterized in the following manner. A loss of signal at a condensate was binned as “detach”. Continuing through the condensate boundary without a change in velocity was binned as “pass”. A “pause” was defined as a diffraction limited stop in processivity for longer than 3 frames (1.5 sec.). These molecules were then binned as pause-pass and pause-detach following the same behaviors as above. “Bidirectional” was binned as molecules that exhibit diffraction limited movement towards a single direction for longer than 1 micron, then reversed direction upon encountering a condensate.

Peak tau intensity for a condensate was derived by first averaging the intensities per pixel over time for a condensate throughout the entire movie. In the event of significant stage drift, the intensities at the time of the events were used. The “Find Peaks” plug-in using default conditions for ImageJ was then used to determine the peak intensity. Background intensity of a non-MT region nearby was then subtracted from this value to determine background subtracted peak intensity.

#### Quantification and Statistical Analysis

All data collected from at least 2 different days with multiple slides per day. Unless otherwise stated, all data was analyzed manually using ImageJ (FIJI). Graphs were created using Graphpad Prism 7.0a and statistical tests were performed using this program. All variances given represent standard deviation. For comparison of DDB behaviors at condensates, the data was collected into a contingency table with four assay conditions and five observable outcomes.

The aggregate analysis of the observed outcome frequencies for the entire table was performed using the Pearson’s chi-squared test and showed significance at very high levels (p < 0.0001). However, some outcome counts were low (below 5) so the analysis was redone using Fisher’s exact test and again significance was extremely high (p < 0.0001). DDB outcomes were compared pairwise with other assay conditions (using data for all outcomes and for just “Pass”/“Pause Pass” pair of outcomes) and significant differences (p < 0.0001) were seen for all comparisons using both tests. In all figures, ^∗^ means p < 0.05, ^∗∗^ means p < 0.01, ^∗∗∗^ means p < 0.001, ^∗∗∗∗^ means p < 0.0001.

**Fig. S1.**
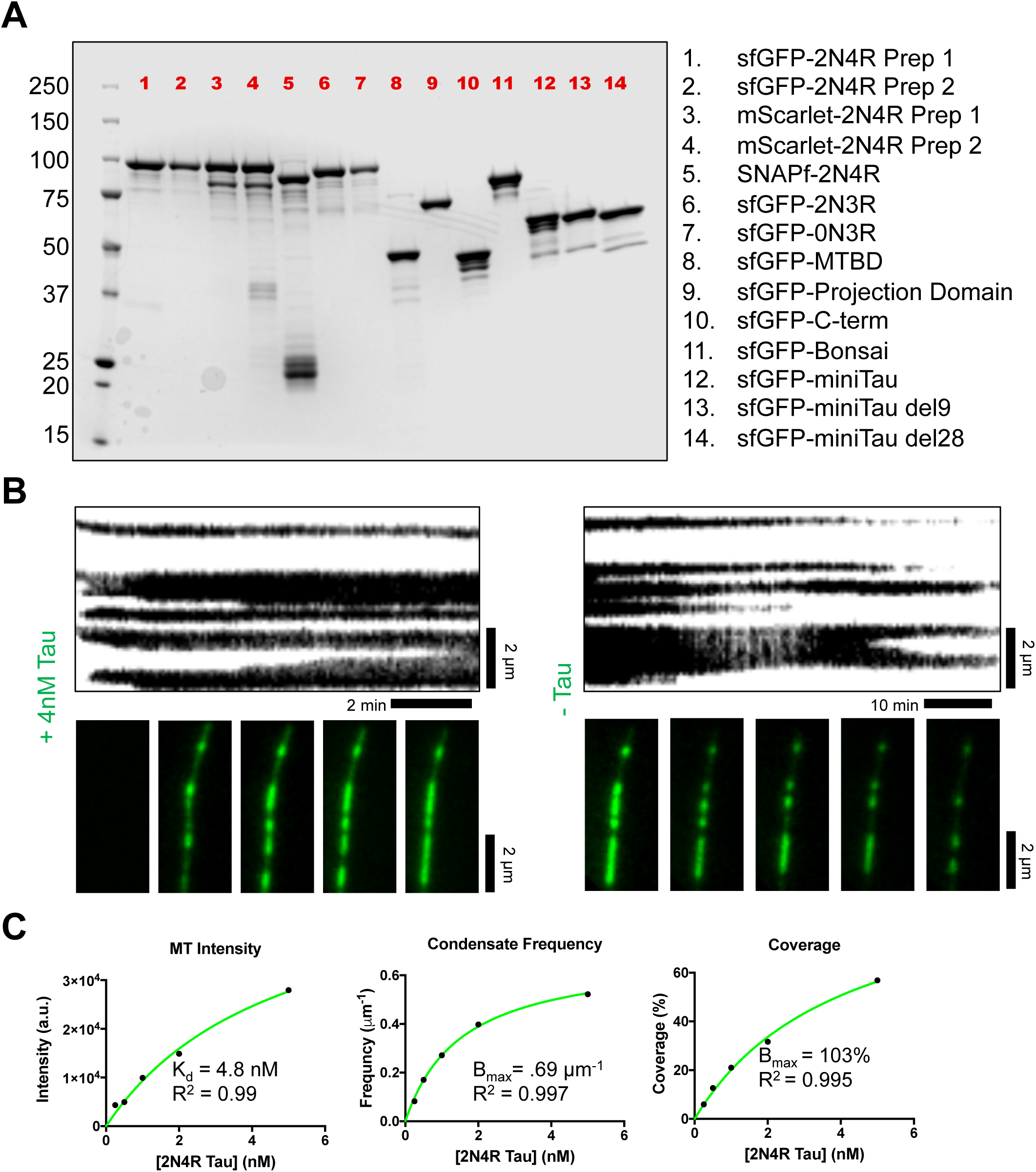
Further Characterization of Tau Condensates. **(A)** SDS-PAGE gel showing proteins used in this study. Protein constructs are listed to the right. **(B)** Kymograph and images of the same MT as GFP-tau is added and removed from the system. Scale bars = 2 μm; 2 min. and 10 min. **(C)** Graphs showing concentration dependence of total MT intensity, frequency, average length, and percent MT coverage of condensates

**Figure S2:**
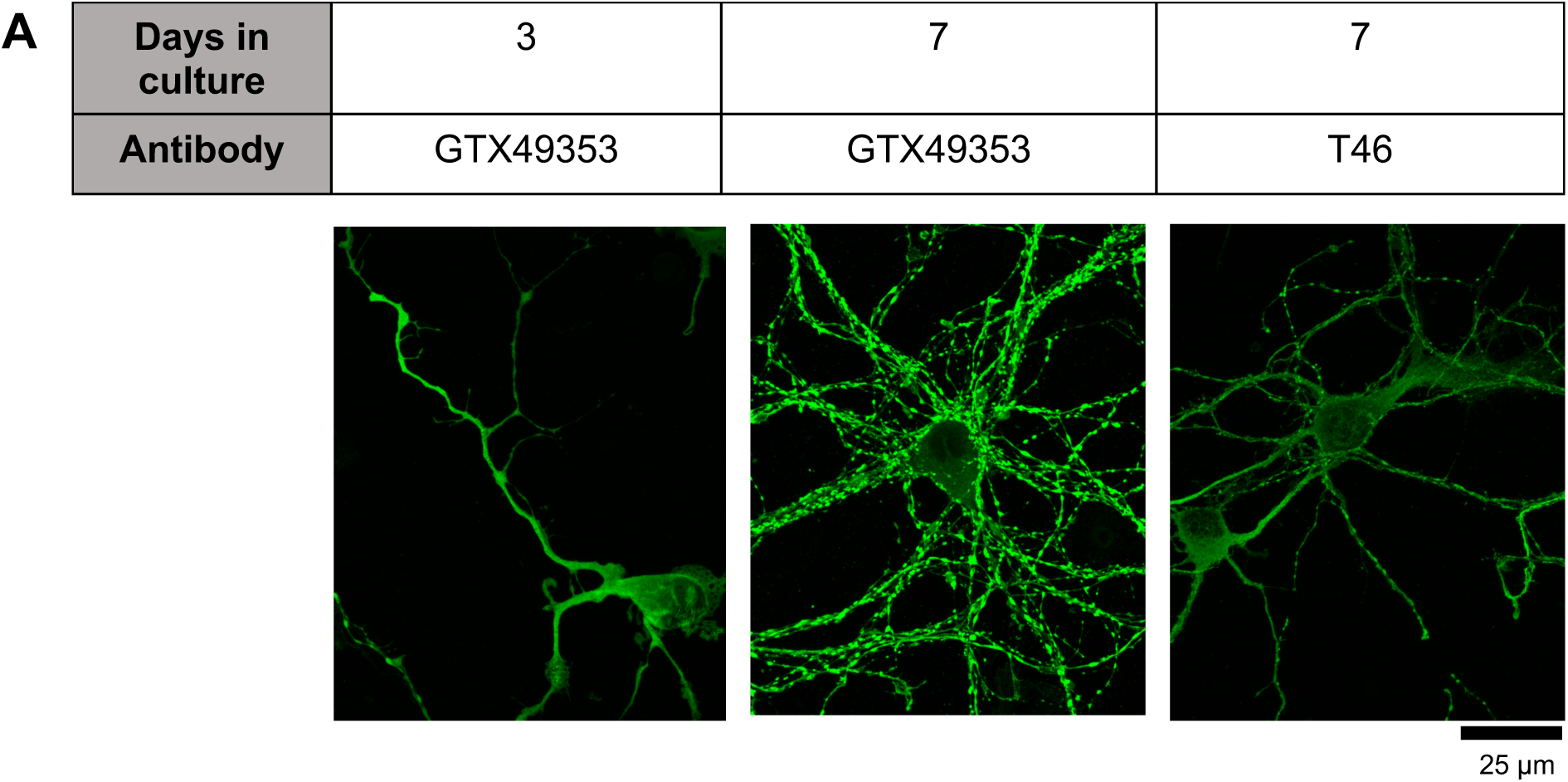
Examples of Total Tau Staining With Different Antibodies. **(A)** Images of mouse hippocampal neurons at different days cultured in vitro. Neurons were immunostained with two different pan-tau antibodies. Scale bar = 25 μm.

**Figure S3:**
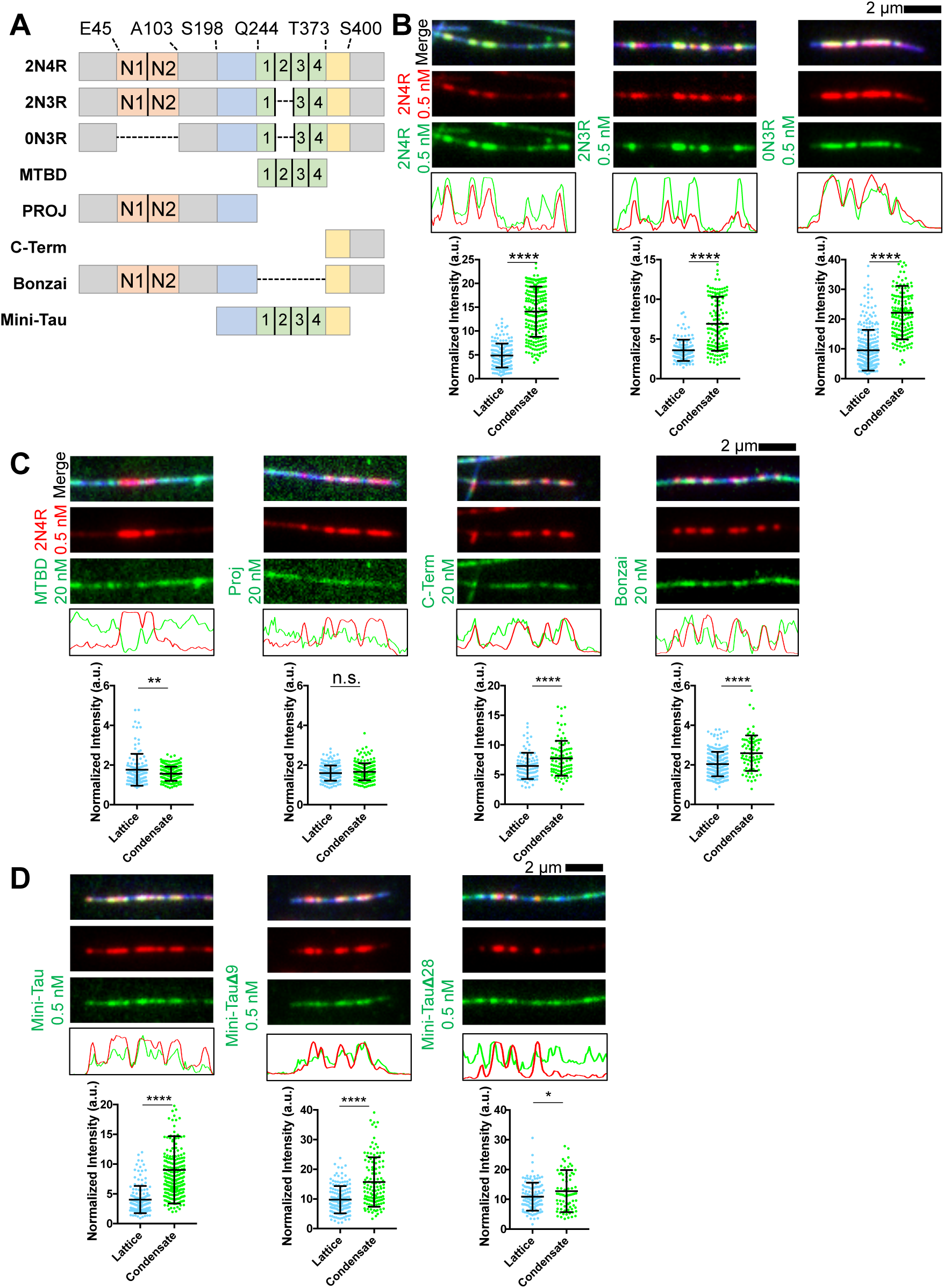
The C-terminal Pseudo-Repeat Region of Tau Licenses the Rest of the Molecule Into Tau Condensates. (A) Schematic of tau isoforms and constructs. Orange boxes: alternatively spliced N-term. inserts. Blue: proline-rich domain. Green: MT binding repeats. Yellow: pseudo-repeat domain. (B) Top: Images of tau isoforms incorporating into full-length tau (2N4R) condensates. Middle: intensity plots of red and green channels. Bottom: tau isoform intensities on the lattice and within condensates. (C) Top: images of constructs incorporating in 2N4R-tau condensates. Middle: intensity plot of red and green channels. Bottom: isoform intensities on the lattice and in condensates. (D) Top: images of mini-tau and pseudo-repeat truncations incorporating into 2N4R-tau condensates. Middle: intensity plot of red and green channels. Bottom: tau isoform intensities on the lattice and in condensates. ^∗^P < 0.05, ^∗∗^ P < 0.01, ^∗∗∗^P < 0.001, ^∗∗∗∗^P < 0.0001. Student’s T-test, one-way ANOVA for multiple comparisons.

**Fig. S4:**
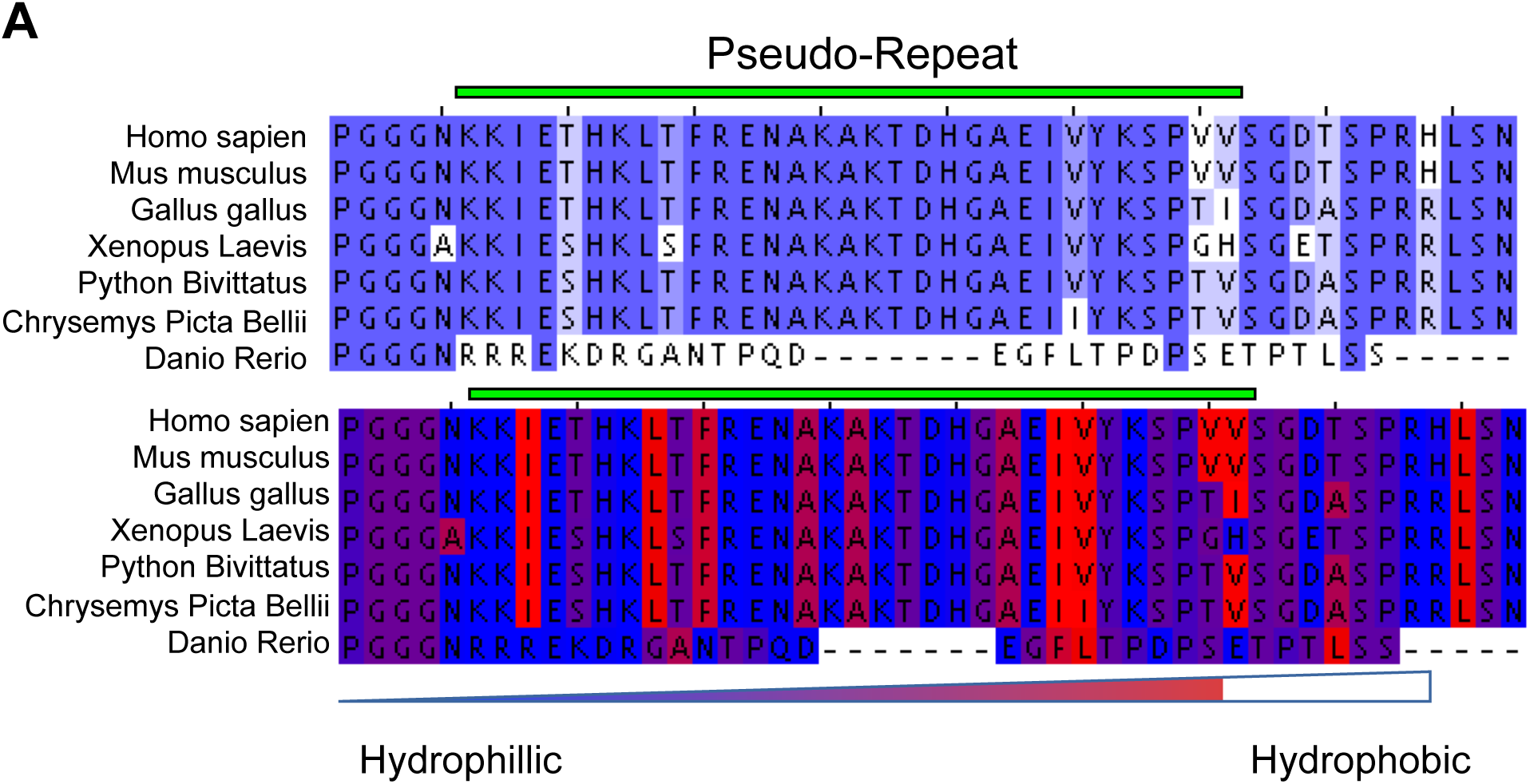
Sequence Alignment of Tau Pseudo-Repeat Region. **(A)** Sequence alignments showing identity conservation (top) and hydrophobicity (bottom) of the tau pseudo-repeat region. The pseudo-repeat is highlighted by green box above.

**Fig. S5:**
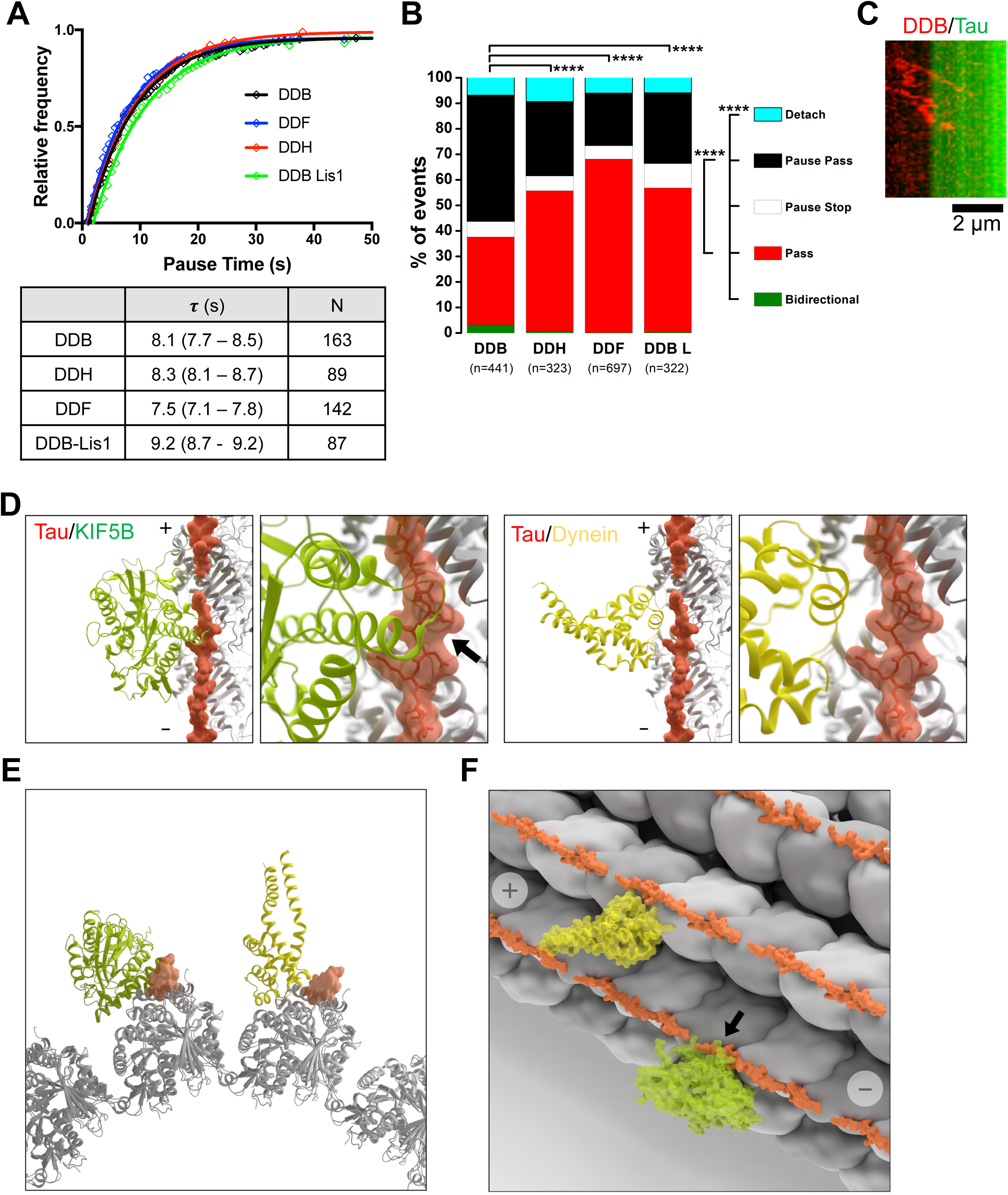
Further Characterization of Tau Condensate Effects on Molecular Motors. **(A)** Cumulative frequency graph and table of pause times for DDX molecules. **(B)** Summary graph of all the behaviors for DDX complexes upon encountering tau condensates. **(C)** Kymograph of a DDB molecule undergoing bidirectional movement when encountering a tau patch. Scale bar = 2 μm; 15 sec. **(D)** Model of MT-binding footprints of dynein, kinesin and tau. (A) Left. Kinesin motor domain (*KIF5B*, pdb: 4HNA; green, *(26)*) footprint at the interface of the tubulin dimer and overlapping tau MT-binding repeats (R2x4, pdb: 6CVN; orange). Right. Dynein motor domain (*DYNC1H1*, pdb: 3JLT; yellow). (E) End-on view of 13 tubulin dimers, shown from the minus (-) end. (F) MT-lattice view. Arrow highlights steric clash between kinesin and tau.

### Supplemental Movie Legends

**Movie S1: Tau forms high density condensates on MTs** Tau condensates (green) forming on a taxol-stabilized MT (blue) when 0.5 nM full-length (2N4R) GFP-tau is added to the chamber. Movie was correct for image drift as described in the Methods. Time in seconds.

**Movie S2: Single molecule spiking experiments reveal tau behavior inside and outside condensates.** Single molecules of 2N4R SNAP-TMR-tau (red, 25 pM) display reduced kinetics within condensates formed with 0.5nM 2N4R GFP-tau (green). Time in seconds.

**Movie S3: Tau condensates form at specific locations on the MT lattice.** Repeated cycles of 1nM full-length (2N4R) GFP-tau (green) and GFP-tau with 8% 1,6-hexanediol reveal ‘hot-spots’ on MTs (blue) for condensate formation. Time in seconds.

**Movie S4: Dynein-Dynactin-BicD2 (DDB) complexes pause at tau condensates.** Single molecules of DDB (red) pause at tau condensates formed with 0.5 nM full-length (2N4R) GFP-tau (green). Time in seconds.

**Movie S5: Tau condensates exclude spastin to protect MTs from severing.** GFP-tau condensates (green) protect regions of MTs (blue) by excluding spastin (red). Note the disappearance of both MT and spastin signal as the MT lattice is destroyed outside of tau condensates. Movie was corrected for image drift as described in the Methods. Time in seconds.

